# Infection Of Rhesus Macaques With O’nyong-nyong Virus UVIR-O804 Recapitulates Key Aspects of Human Clinical Disease

**DOI:** 10.1101/2025.11.20.689439

**Authors:** Hannah K. Jaeger, Michael Denton, Takeshi F. Andoh, Craig N. Kreklywich, Lina Gao, Lydia J. Pung, Zachary J. Streblow, Ann McMonigal, Karina Ray, Brayden Graves, Magdalene M. Streblow, Aaron M. Barber-Axthelm, Gavin Zilverberg, Margaret Terry, Suzanne S. Fei, Glenn Hogan, David C. Schultz, Sara Cherry, Michael K. Axthelm, Caralyn S. Labriola, Mark Heise, Daniel N. Streblow

## Abstract

1.

O’nyong-nyong virus (ONNV) is a mosquito-borne alphavirus first isolated in Uganda in 1959. Since its discovery, ONNV has caused several outbreaks in Africa, manifesting clinically as fever, rash, and joint/muscle pain lasting months. Currently, we have a limited understanding of ONNV infection and disease in relevant animal models, which restricts the evaluation of vaccines and therapeutics. In 1967, Binn et al. reported that infection of rhesus macaques (RMs) with ONNV failed to induce viremia in two animals. This may be attributed to the potential attenuation of the virus through extensive passaging. To mitigate this issue, we constructed an infectious clone from the sequence of ONNV-UVRI0804 (ONNV_0804_), a 2017 clinical isolate from a febrile patient in Uganda. This strain demonstrated high pathogenicity in immunocompetent mice, resulting in an earlier and more severe onset of disease and significantly higher viremia compared to a highly passaged control strain ONNV_UgMP30_. In the current study, three male and three female rhesus macaques were subcutaneously inoculated with ONNV_0804_. All animals became viremic at 2 days post inoculation (dpi). Both classical and nonclassical monocytes were activated (CD169+), peaking at 3 dpi, which corresponded with the peak of viremia. Additionally, CD4+ and CD8+ effector memory T cells and memory B cells began proliferating in peripheral blood by day 7, peaking at day 10, which also corresponded to the timing of neutralizing antibody development, indicating a robust adaptive immune response to ONNV_0804_. Finally, key clinical disease manifestations were recapitulated, including lymphadenopathy and histological features of early-stage arthritis. Taken together, rhesus macaque infection with ONNV_0804_ clinical isolate is a promising model for investigating immune responses to alphaviruses and evaluating vaccines to protect against future epidemics.

**Author summary:** O’nyong-nyong virus (ONNV) is a mosquito-transmitted alphavirus that causes fever, rash, and prolonged joint and muscle pain, similar to chikungunya virus and other arthritogenic alphaviruses. Despite its capacity to cause outbreaks in Africa, ONNV remains understudied, and there are currently no approved vaccines or therapeutics to prevent or treat infection and disease. A major barrier to advancing ONNV research has been the lack of suitable animal models to study the virus and investigate host immune responses. We engineered an ONNV infectious clone of a recent clinical isolate sequenced from a patient in Uganda (ONNV_0804_) that causes robust infection and disease in immunocompetent mice. In the current study, we provide data demonstrating that this contemporary ONNV strain infects rhesus macaques. Notably, rhesus macaques developed detectable viremia, rash, lymphadenopathy, joint and muscle inflammation, and strong innate and adaptive immune responses following subcutaneous ONNV_0804_ infection. These findings suggest that ONNV_0804_ infection in macaques is a promising model for studying ONNV pathogenesis and immunity. This model will be instrumental for evaluating future vaccine and therapeutic candidates aimed at preventing ONNV infection and related viral disease.

## 3. Introduction

O’nyong-nyong virus (ONNV) is a mosquito-transmitted alphavirus first identified in Uganda in the 1959 outbreak (1). ONNV then reemerged to cause two additional major outbreaks in Uganda in 1996 and Chad in 2004 (2, 3, 4). Other sporadic cases have occurred across Africa but have not reached outbreak levels. Serology studies show ONNV is an emerging and reemerging neglected tropical disease; however, clinical surveillance for active infections remains limited due to clinically indistinguishable symptoms with other arboviruses (5). ONNV has the potential for enzootic and urban transmission cycles due to its ability to infect both *Anopheles* and *Aedes* mosquitoes (6, 7, 8), which is a concern for spread into new regions. The ability to infect multiple mosquito species also increases the risk of zoonotic spillover, leading to potential urban outbreaks. Further, vector-borne diseases account for 700,000 deaths annually (9, 10) with close to 3.9 billion people living in subtropical and tropical climates where arboviruses circulate, creating a global health threat that disproportionately affects communities with limited socioeconomic resources (11).

In humans, ONNV infection manifests with fever, rash, and debilitating joint and muscle pain that can persist well past the timing of detectable viremia (12, 13, 14). The clinical course of ONNV is similar to that of chikungunya virus (CHIKV) and begins with a 4–7 day incubation followed by acute disease symptoms that last one to two weeks. Some individuals develop chronic or persistent joint pain and fatigue that may last from weeks to years (15, 16, 17, 18, 19). ONNV differs from related arthritogenic alphaviruses, including CHIKV (20) and Mayaro (MAYV) (21), in that acute joint edema is not typically observed (12). Instead, lymphadenopathy has been noted as a distinctive manifestation of ONNV infection (1, 2, 22). The clinical similarity to other arboviral diseases such as chikungunya, dengue, and Zika viruses often leads to diagnostic challenges, underreporting, and frequent disease misclassification. Even neutralization assays, the current serological gold standard, often cross-react with closely related alphaviruses, including CHIKV (23). At present, there are no licensed vaccines or specific antiviral treatments for ONNV, reflecting the broader challenge of neglected tropical diseases.

The potential for ONNV to extend beyond its traditional geographic range, combined with the lack of preventative or therapeutic options, underscores the urgent need for research on effective vaccines and antivirals. A major obstacle to advancing this research is the paucity of a well-validated animal model. Mouse models, including those using immunocompromised strains, have provided valuable insights into alphavirus replication, immune responses, and pathogenesis. Although ONNV strain-specific differences in clinical manifestation have not yet been identified, differences in pathogenicity and immune responses have been observed in mice (24, 25). However, mice do not always fully recapitulate the human clinical disease course, particularly the chronic arthralgia and joint pathology that define arthritogenic alphavirus infections. The short lifespan and limited complexity of the murine immune system restrict the ability to study persistent or relapsing joint inflammation, or the durability of immune responses, all critical components of human disease.

Nonhuman primates (NHP) share close immunological, genetic, and physiological similarities with humans, making them suitable for modeling persistent musculoskeletal disease, evaluating the kinetics of innate and adaptive immunity, and defining correlates of vaccine-mediated protection (26, 27, 28, 29, 30). Previous attempts to model ONNV pathogenesis in rhesus macaques failed to produce viremia, although the virus passage history was not described and the virus may have been attenuated, which limits the interpretation of those results (31). Serial laboratory passaging of alphaviruses is known to generate mutations that reduce virulence *in vivo* (32, 33). Thus, our understanding of ONNV pathogenesis and host responses in a relevant model remains incomplete, and the field lacks a robust nonhuman primate (NHP) model to evaluate vaccine or therapeutic efficacy.

NHP models of related arthritogenic viruses like CHIKV and MAYV infection have already demonstrated their value. Established infection models in cynomolgus macaques (*Macaca fascicularis*) (34, 35) and rhesus macaques (*Macaca mulatta*; RM) across adult, aged, and pregnant cohorts (31, 36, 37, 38) recapitulate many aspects of human disease and have provided a critical platform for evaluating vaccines and monoclonal antibody therapies (39, 40, 41). These models have established that NHPs can serve as translationally relevant systems for arthritogenic alphavirus research, yet important questions remain unanswered regarding ONNV viral tissue tropism, persistence, strain-specific differences in pathogenicity, and the durability of innate and adaptive immune responses in a relevant model. To address these gaps, we engineered an ONNV_0804_ infectious clone that was derived from a sequence obtained from a patient in Uganda in 2017 (42). This clone is virulent in mice, causing robust acute and persistent infection accompanied by arthritogenic disease, and provides a foundation for the development of a relevant NHP model (24). To our knowledge, this study represents the first productive ONNV infection that recapitulates human arthritogenic disease in NHP. This model provides an essential platform to define correlates of protection, investigate mechanisms of viral persistence and musculoskeletal pathology, and accelerate the development of vaccines and therapeutics against this neglected tropical virus.

## 4. Results

### 4.1 Kinetics of ONNV replication in rhesus macaques

To characterize ONNV disease pathogenesis in nonhuman primates, six rhesus macaques (RM); three females (ages 8, 11, 12 years old) and three males (ages 4, 4, and 5 years old) were used in this study (**Figure 1A**). Similar to previous models of arboviruses (26, 35, 43), RMs were inoculated subcutaneously (s.q.) in the hands and arms bilaterally at five sites per side (100 μL per site) in an attempt to mimic mosquito bites with a total infectious dose of 1 × 10^5^ plaque forming units (PFU) of ONNV_0804_. Clinical assessments, peripheral blood, and urine samples were collected at 0, 1, 2, 3, 5, 7, 10, and 14 days post-inoculation (dpi) **(Figure 1A)**. RM were humanely euthanized at 14 dpi for tissue collection to determine viral biodistribution and immunological assays to understand relevant host responses to viral infection, which was chosen to better capture the development of adaptive immune responses (26, 35).

**Figure 1.**
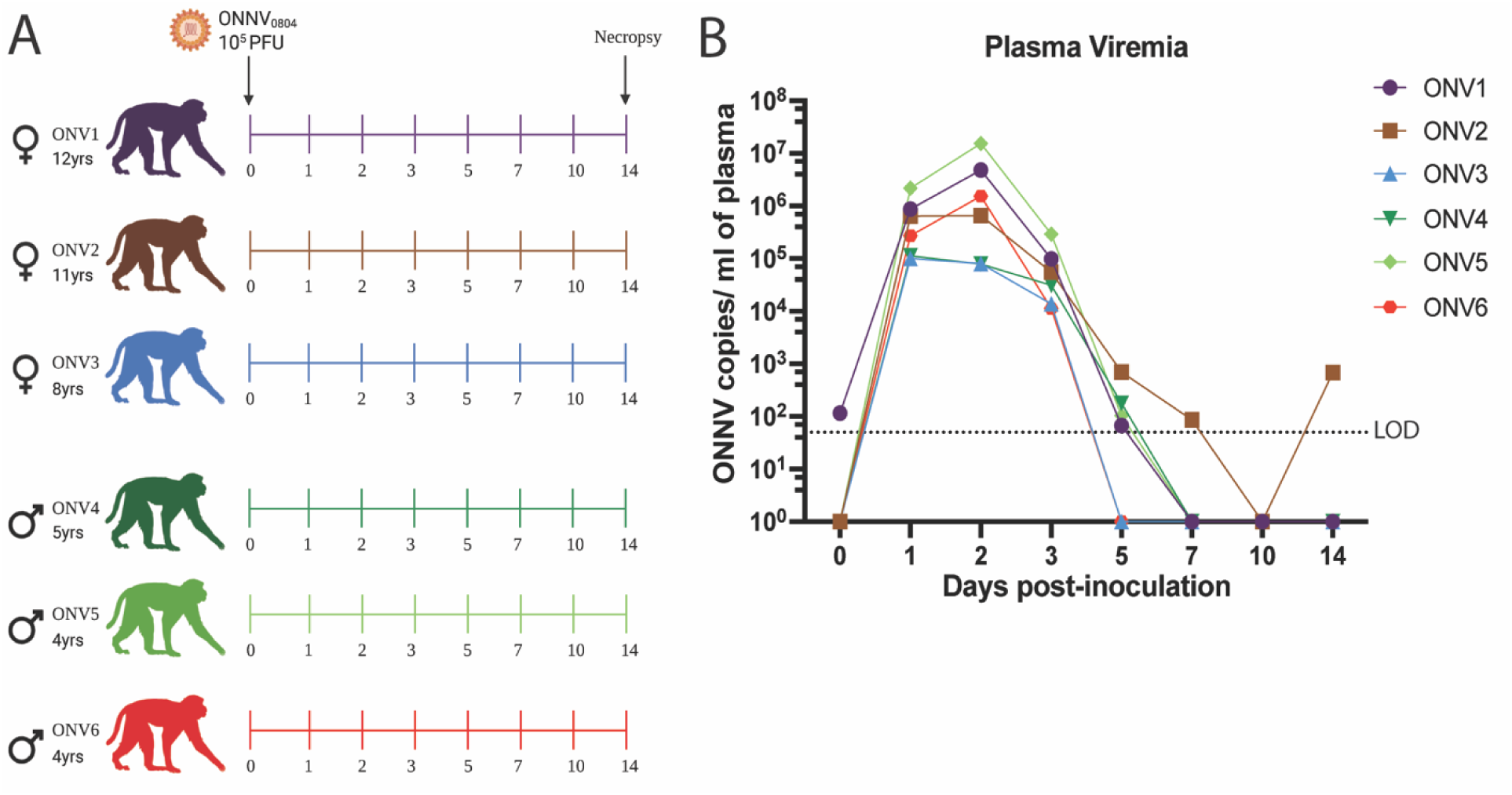
Study overview of ONNV infection of NHPs. **(A)** Schematic summarizing the ONNV rhesus macaque infection study. Animals were inoculated with 1 x 10^5^ plaque forming units (PFU) of ONNV_0804_ administered s.q. and spread evenly in both arms and hands. Blood was drawn for PBMC and plasma isolation as well as complete blood count (CBC) 0–5, 7, 10, and 14 dpi. Animals were humanely euthanized at 14 dpi and extensive lymphoid, muscles, joints, nerves, lobes of the brain, heart, major organs, and reproductive tissues were harvested. (**B**) Viral RNA detection in plasma using qRT-PCR with data representative of three independent replicates. The LOD was 50 copies ONNV RNA per mL of plasma with undetectable samples graphed as 0 copies vRNA/mL plasma.

Viral RNA (vRNA) detected in plasma following infection peaked between 1-2 dpi, with a mean peak viremia of 9.8 × 10⁵ copies/mL **(Figure 1B)**. Repeated-measures one-way ANOVA (Geisser–Greenhouse corrected) revealed significant differences across time (p < 0.0001). Post-hoc Sidak’s multiple-comparisons test showed that mean plasma viral loads at 1–3 dpi were significantly higher than baseline (p < 0.0001 for all), whereas by 5 dpi, viremia declined and was no longer significantly different from baseline (p = 0.1453). One animal, ONV2, had detectable viremia at 14 dpi at low levels, not uncommon for mosquito-borne viruses to cause secondary viremia in the host. Based on peak levels of viremia, the animals naturally divided into two groups, with ONV1, ONV2, ONV5, and ONV6 having the highest viremia peaks, and the highest individual peak viremia observed being 1.54 × 10⁷ copies/mL in ONV5. The second group, consisting of ONV3 and ONV4, had an approximate 10-fold lower vRNA detected at the peak of viremia, and these animals also experienced a shorter viremic duration. Area under the curve (AUC) was compared between male and female animals using an unpaired t-test (Mann–Whitney test). No significant difference was detected (p = 0.52), consistent with similar magnitude and duration of viremia between sexes. One animal (ONV1) exhibited detectable viral shedding in urine at 1 and 2 dpi; although low in magnitude (1.4 × 10³ copies/mL at peak). This finding corresponded temporally with its plasma viremia peak of 1.54 × 10⁷ copies/mL at 2 dpi, which was the highest for any of the animals **(S Figure 1)**. This demonstrates moderate individual variability but supports this model’s reproducibility in consistent and comparable viremia across animals.

### 4.2 ONNV-induced tissue tropism and immunopathology

To evaluate changes in clinical symptoms of infection, body temperature, weight, skin, and joint changes were monitored following infection. Temperature was assessed every morning during blood draws, but no significant changes or fever were observed in any of the animals **(S Figure 2A)**. This finding is postulated to reflect the timing of readings, as fever spikes in febrile illnesses often occur at night (44). None of the six animals experienced significant weight loss over the duration of the 14-day infection study, although ONV4 did experience 5% loss of body weight post-challenge **(S Figure 2B)**. No other signs of distress were noted in any of the animals. Complete blood counts of each macaque across the study timepoints revealed few remarkable changes, but rather followed typical infection dynamics **(S Figure 3)**. Neutrophils slightly spiked at 1 dpi and decreased afterward, while the frequency of monocytes slightly increased at 2 dpi and remained elevated **(S Figure 3A, B)**. Lymphocytes appeared to increase in frequency at later times post-infection in most animals **(S Figure 3C)**.

Two animals developed bilateral dermal lesions on the arms at 7 dpi, consisting of edema of the dorsal wrists and hands (ONV5) and a maculopapular rash affecting the arms and shoulders (ONV6) **(Figure 2A-C)**. ONV4 had ventral seborrheic dermatitis prior to infection with no changes noted throughout the study and developed a small superior lip lesion. By 14 dpi, all animals except ONV1 and ONV4 presented with dermal pathology of the upper limbs, including hyperkeratosis, erythema, and edema of the dorsal hands and wrists, axillary macular rash, and maculopapular rash **(Figure 2A,D, S Figure 4A)**. Pruritus was not observed in any animal. Bacterial culture of the rashes revealed normal background dermatologic flora, and histologic screening for other etiologic agents, such as measles virus, was negative. On histologic examination, the skin lesions displayed varying degrees of increased perivascular lymphocytic aggregates, lymphocytic to lymphoplasmacytic dermatitis, acanthosis, and hyperkeratosis with severity mirroring that of the gross findings **(Figure 2B,C, S Figure 4 B-D)**. The most affected areas had lymphoplasmacytic inflammation extending to the follicles and adnexal structures when compared to control skin taken from abdomen. All skin lesions and rashes appearing after inoculation that were tested for ONNV vRNA RT-qPCR were found to be positive **(Figure 2G, I, S Table 1)**. Three animals also developed lymphadenopathy of one or multiple peripheral lymph node groups (ONV3, ONV5, ONV6), a key clinical symptom of ONNV in humans, which corresponded microscopically to lymphofollicular hyperplasia **(Figure 2 D-F)**. Taken together, these data show the rhesus macaque model recapitulates key pathologic features of ONNV infection in humans.

**Figure 2.**
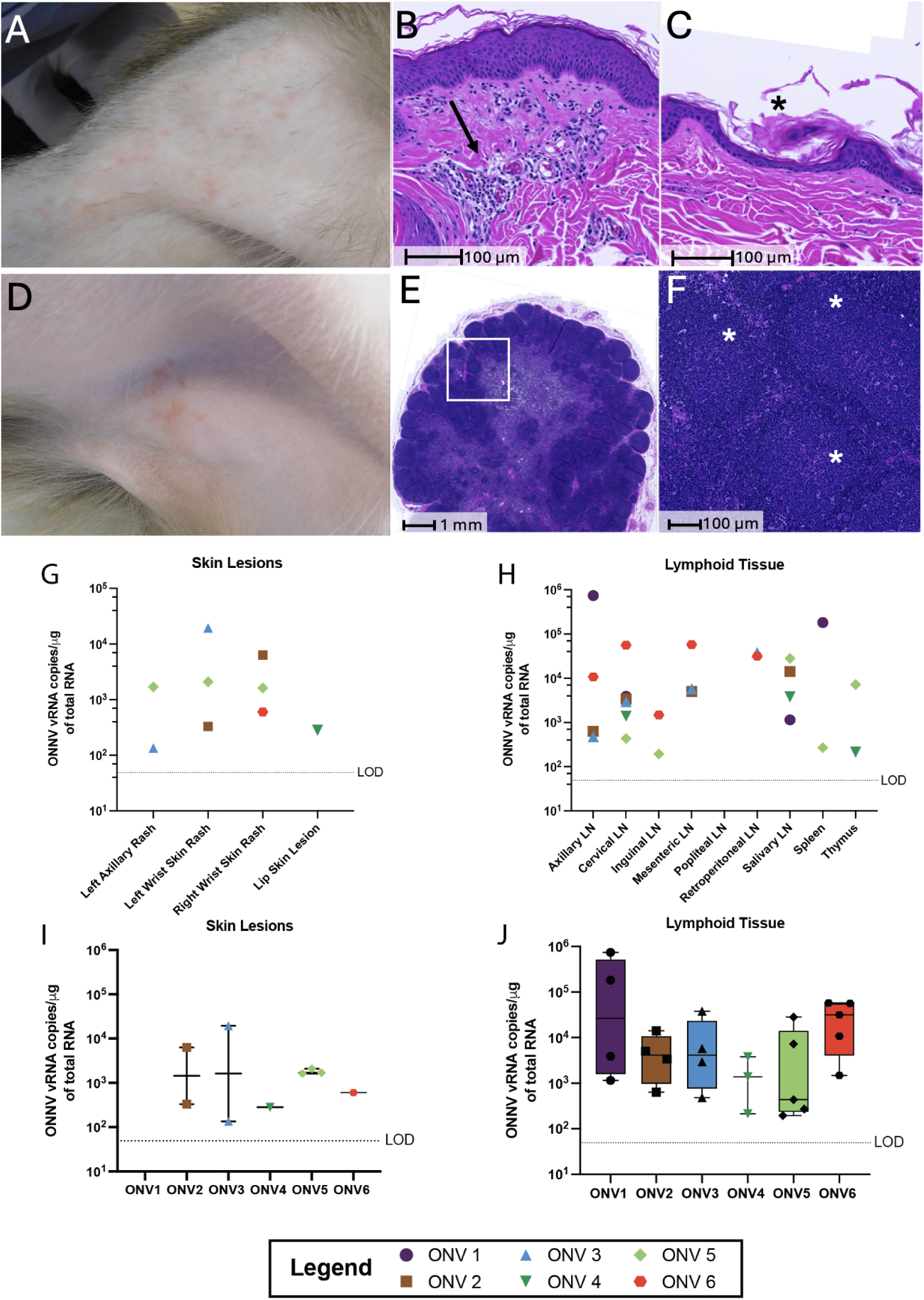
Dermatologic pathology and viral burden in ONNV-infected rhesus macaques. Gross images of representative dermatologic manifestations, including maculopapular rash on the shoulder and brachium **(*A*)** and axillary rash with lymphadenopathy **(*D*)**. At 14 dpi, skin tissue sections with grossly visible lesions and associated lymphoid tissue were fixed, paraffin-embedded, sectioned at 5 μm, and stained with hematoxylin and eosin (H&E) for histological evaluation. (***B,C*)** Sections of rash displaying multifocal acanthosis, mild hyperkeratosis, perivascular lymphocytic inflammation and superficial dermal edema in the dermis. Mild lymphofollicular hyperplasia (white asterisk) was present on histologic examination of the axillary lymph node at low **(*E*)** and high **(*F*)** magnification. Viral loads (from **S. Table 1**) were quantified using one-step RT-qPCR on the RNA isolated from 14 dpi necropsy samples. The limit of detection (LOD) was ∼50 copies of ONNV RNA. Graphs represent tissues with at least one animal positive for viral detection and are separated into five tissue types. Skin **(*G*)** and lymphoid tissues **(*H*)** are shown, with summary graphs **(*I,J*)** presented below each tissue group.

Each of the animals infected with ONNV exhibited a spectrum of joint pathology, most common and severely manifesting within the upper limbs. At necropsy, minimal to mild effusions were present in the wrists plus or minus elbow and finger joints of three animals (ONV2, ONV3, ONV6). Slides from the elbows, wrists, combined metacarpophalangeal and carpal interphalangeal, stifles, ankles, and combined metatarsophalangeal and tarsal interphalangeal joints were evaluated by T.F.A., M.K.A. and C.S.L. The periarticular tissues of two or multiple joints, in all animals, showed a range from minimal perivascular inflammatory cell aggregates to moderate lymphoplasmacytic and neutrophilic inflammation (**Figure 3A-D)**. Across the entire cohort, the most prevalent changes were found in the periarticular adipose tissue and loose connective tissue, with few sub-synovial foci and one perivascular lymphocytic aggregate within the periosteum. Similar to what we recently reported in the MAYV NHP model and ONNV challenged C57BL/6 mice (24, 26), evidence of periarthritis and sub-synovial steatitis after inoculation of rhesus macaques with ONNV was clearly demonstrated **(S. Figure 5A-C)**. To quantify these findings, inflammation was scored based upon a histologic grading rubric for both dermatologic findings and joint pathology (**S Table 2**). All six animals had scores between 1 to 8 within at least one joint type (**Figure 3E, S Table 3)**. While there was no clear trend relating to the magnitude of plasma viremia, the histologic scores of the wrists, fingers, and elbows reflected the gross presentations of rash and edema in nearby joints.

**Figure 3.**
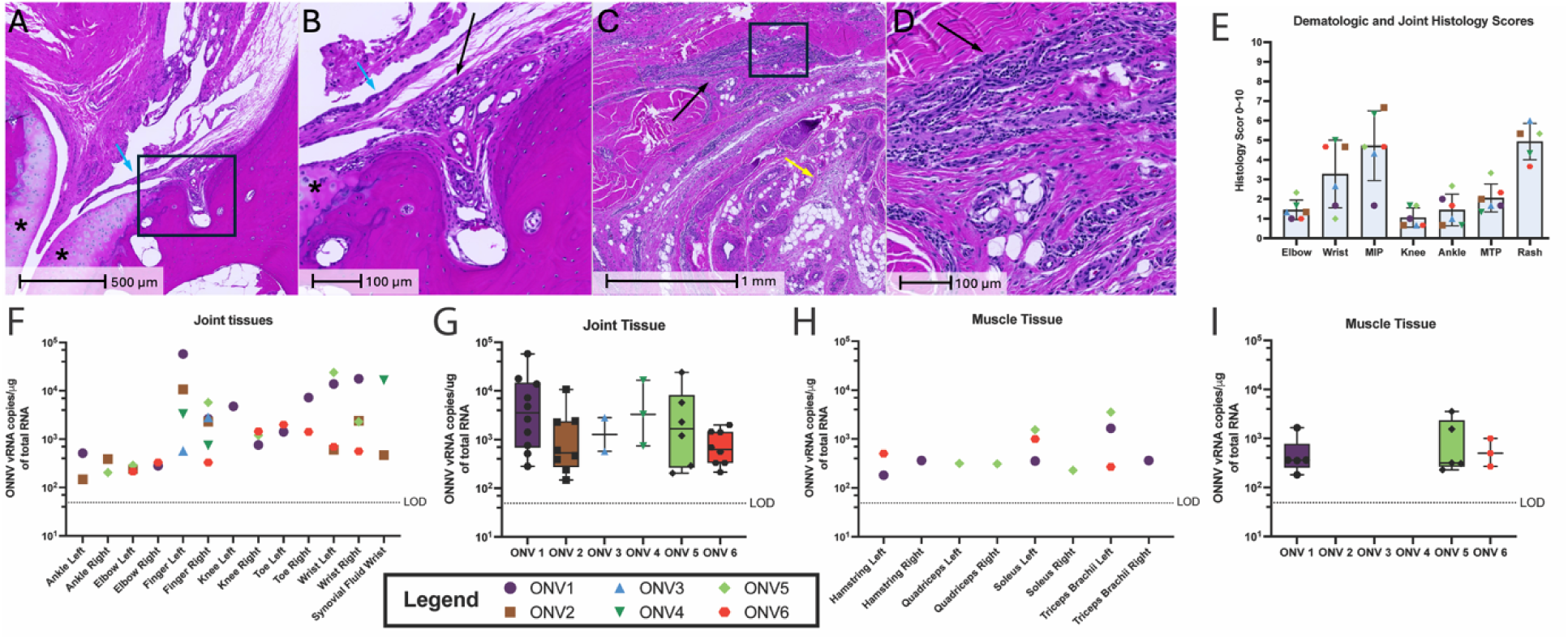
Histopathological and virological assessment of ONNV-infected joints and muscles. Representative images of finger tissue at 14 dpi stained with H&E displaying joint inflammation. Histologic analysis of a carpal interphalangeal joint from ONV5 at low (***A***) and high (***B***) magnification revealed sub-synovial lymphocytic inflammation (black arrow) with minimal focal hypertrophy of the overlying synovium (blue arrow). The articular surfaces (black asterisk) were unaffected. Lymphoplasmacytic and neutrophilic inflammation evident at low (***C***) and high (***D***) magnification extended within the tendon (black arrows) and adipose tissue (yellow arrow) of ONV4. (***E*)** Combined histology scores averaged across three scores for selected tissues. Sections were evaluated for perivascular lymphoid aggregates, inflammation, and synovial hypertrophy or hyperplasia. Severity scores were assigned on a scale of 0-10, as described in **S Table 2**, with 0 indicating the absence of any changes, 1-2 scarce, 3-4 minimal, 5-6 mild, 7-8 moderate, and 9-10 severe. Viral loads (from **S. Table 1**) were quantified using one-step RT-qPCR. The limit of detection (LOD) was 50 copies of ONNV RNA. Graphs represent tissues with at least one animal positive for viral detection and are separated into five tissue types. Joint and muscle tissues were summarized separately **(*G, I*)**.

Since all animals experienced joint pathology, we determined the tissue distribution of ONNV vRNA in the joints and muscles. High levels of vRNA were detected in the right and left muscle and joint tissues, including ankles, toes, elbows, fingers, and wrists for all six animals, indicating that the virus disseminated effectively throughout the body **(Figure 3F-I, S Table 1).** Synovial fluid was successfully isolated from the wrists of ONV2 and ONV4; however, this was not feasible for all animals, which limits interpretability **(S Table 1)**. No virus was isolated from the elbow or ankle synovium at necropsy (14 dpi). Summary graphs **(Figure 3G,I)** show the breadth and magnitude in tissue subsets and highlight that viral RNA levels detected post-inoculation correlate with the extent of tissue distribution. For example, ONV3 and ONV4 had the lowest peak and duration of viremia, which corresponds with fewer joints with detectable ONNV vRNA, and these animals also did not have any vRNA-positive muscle tissues. These data provide valuable insights into ONNV tissue tropism, replication, distribution, and persistence, which align with findings from previous mouse studies (24, 25).

### 4.3 ONNV disseminates to the central and peripheral nervous system tissues

Studies with MAYV, a related alphavirus, detected histologic changes and vRNA within NHP major organ systems (26). In the current ONNV study, major organ systems were screened for pathology by histology; however, only minor or mild inflammatory cell pathologies were present in some animals. In the heart, minor focal aggregates of lymphocytes were observed in animals ONV1 and ONV4 (**Figure 4A**). ONV5 had mild hyperplasia of the bronchus-associated lymphoid tissue (**Figure 4B**). Renal findings within ONV1, ONV4, and ONV5 were minimal perivascular lymphocytic aggregates, mild interstitial lymphocytic infiltrates in the cortex, and tertiary lymphoid structure formation in the pelvis (**Figure 4C**). These findings are within the realm of background findings in macaques (45), though they may spur future studies into co-localization of virus to pathology, as the heart of ONV1 and lungs and kidneys of ONV5 had relatively higher vRNA levels in the major organ systems (**S Table 1**).

**Figure 4.**
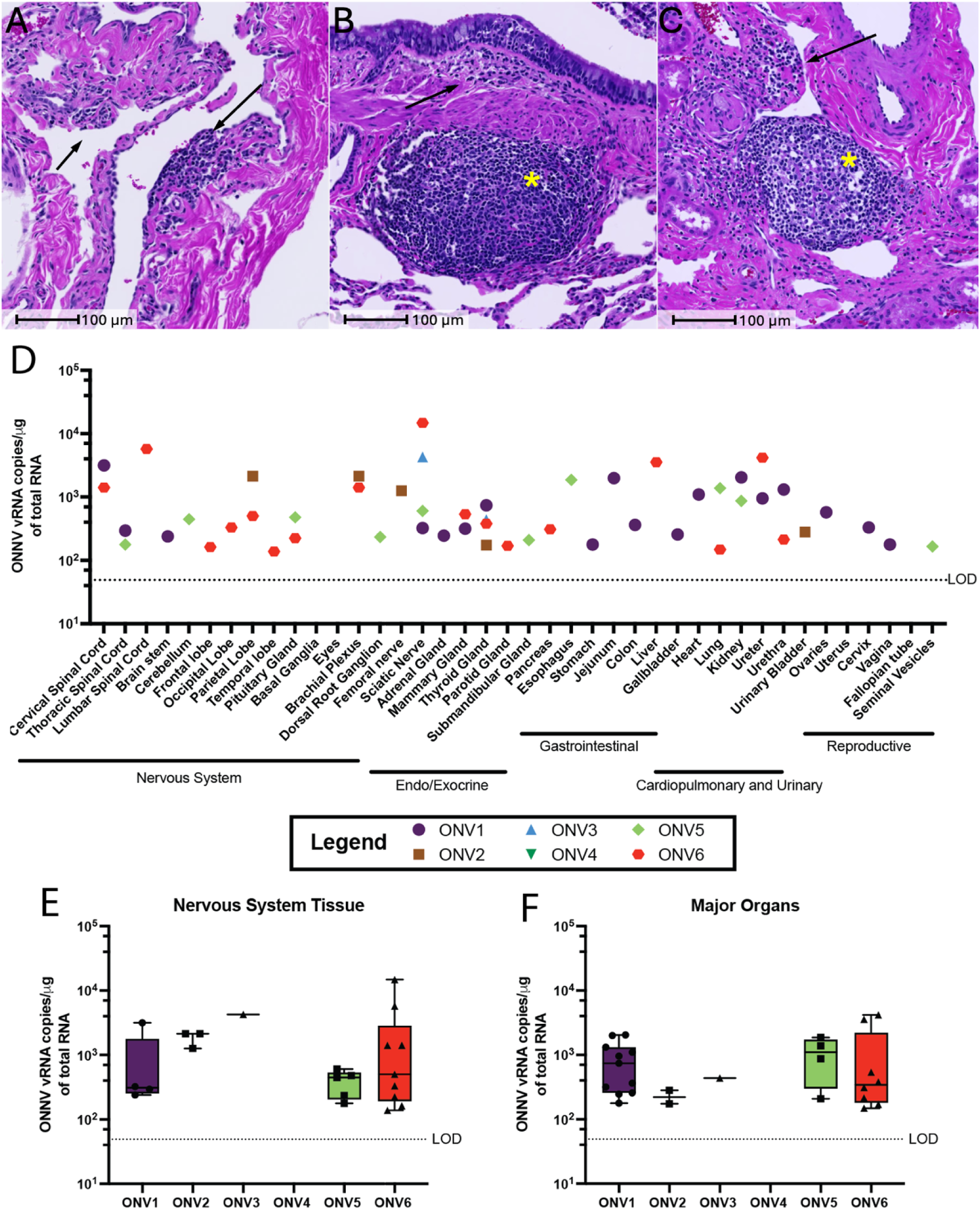
Lymphocytic inflammation in heart, lung and kidney of ONNV_0804_ infected rhesus macaques. Representative lymphoid findings on H&E-stained sections from the major organs. **(*A*),** Minor perivascular aggregates of lymphocytes (black arrows) within the connective tissue adjacent to the aortic trunk in a section of heart tissue from ONV1. **(*B*),** Section of lungs from ONV5 showing minor lymphocytic infiltration into the bronchial epithelium and submucosa (black arrow) with mild hyperplasia of bronchus-associated lymphoid tissue (yellow asterisk). **(*C*),** Minor perivascular lymphoid aggregates (black arrow) and tertiary lymphoid structure formation (yellow asterisk) in the renal pelvis of ONV5. **(*D*),** Nervous system (CNS & PNS), gastrointestinal, cardiopulmonary and urinary, and reproductive were quantified with summary graphs for nervous system tissue and major organs **(E,F)**.

Alphaviruses are divided into encephalitic and arthritogenic viruses based on their primary disease manifestations. Encephalitic alphaviruses, including Eastern Equine Encephalitis Virus (EEEV), Western Equine Encephalitis Virus (WEEV), and Venezuelan Equine Encephalitis Virus (VEEV), cause viral-induced encephalitis and neurologic disease within 10-14 days post-infection that can progress to death. Arthritogenic alphaviruses, including CHIKV, MAYV, and Ross River virus, typically do not cause encephalitis or other neurological complications. Emerging evidence further indicates that these arthritogenic viruses can infect nervous system tissues. However, neurological complications can happen in those 65 years and older, neonates, or immunocompromised individuals who become infected with these arthritogenic viruses (46). To assess whether ONNV crosses the blood–brain barrier in NHPs, we first measured vRNA in the cerebrospinal fluid, where no animals showed detectable viral RNA. In contrast, vRNA was present in multiple brain lobes and major central nervous system regions across all six animals. The basal ganglia and trigeminal ganglion were the only sampled nervous system tissues in which viral RNA was not detected for any animal **(Figure 4D, S Table 1)**. Although we detected ONNV vRNA in many nervous system tissues, we did not observe evidence of neurological disease in any of the macaques when evaluated clinically or histologically.

The major organ systems were also tested for the presence of vRNA. In the gastrointestinal system, the duodenum and ileum were the only tissues that were undetectable for vRNA in any of the animals **(Figure 4D, S Table 1)**, which is consistent with the MAYV NHP model (26). ONNV RNA was detectable in the heart, kidney, urethra, and ureter of ONV1, and the lungs and kidney of ONV5. Viral RNA was detected at low levels in the female reproductive tissues and was also found in the seminal vesicles of ONV5 **(Figure 4D, S Table 1)**. Of note, ONV2 and ONV3 had a previous ovariectomy; therefore, the full extent of dissemination seen for reproductive tissues could not be completed. We compared the pooled vRNA levels in nervous system tissues and major organs, and there were no significant differences detected between animals **(Figure 4D, E)**.

To assess the relationship between systemic viremia and tissue distribution, we ran a correlation analysis between peak viremia and percent tissue distribution. A positive correlation was observed between peak viremia and the proportion of tissues positive for viral RNA (Spearman r = 0.77, p = 0.10), indicating that animals with higher viremia tended to have broader tissue dissemination, although this trend did not reach statistical significance. Together, the distribution of vRNA and associated histopathologic changes indicate that ONNV infection in rhesus macaques is systemic and targets multiple organ systems before resolution.

### 4.4 Transcriptomic analysis reveals host responses that are conserved between ONNV and MAYV

We next profiled longitudinal PBMC transcriptomes by RNA-seq to capture early systemic responses to infection. We focused our analysis on samples obtained during peak viremia (2 dpi v 0 dpi). Differential expression (DE) analysis highlights the strong interferon response and antiviral responses with top upregulated genes (FDR p<0.05, FC (fold change) >2), including IFI27, ISG15, MYO7B, HERC5, IFI6, RSAD2, IFIT1, APOBEC3A, OAS1, and FIT3 **(Figure 5A)**. These genes are upregulated significantly when compared to baseline at 0 dpi and remain elevated to a lesser extent at 3 dpi before returning to baseline at 10 dpi. Ingenuity pathway analysis (IPA) was used to further understand the biological mechanisms induced by infection. Top pathways included type I/II interferon responses with downstream targets such as ISG15, IFI6, IFI27, OAS1, MX1, IFIT2, IFIT3, IFIT5, and IFNG-inducible TRIMs **(Figure 5B, C)**. Pattern recognition and innate antiviral responses were also prominent with genes that trigger immune responses (RIGI, TLR7, and MAVs) and several genes involved in antiviral responses (EIF2AK2, IRF7, and STAT1). A similar transcriptional response is observed at 3 dpi but is largely gone by 5 dpi **(Figure 5D)**. These findings suggest innate immune modulation plays an important role in regulating antiviral immunity and early control of the virus following ONNV infection.

**Figure 5.**
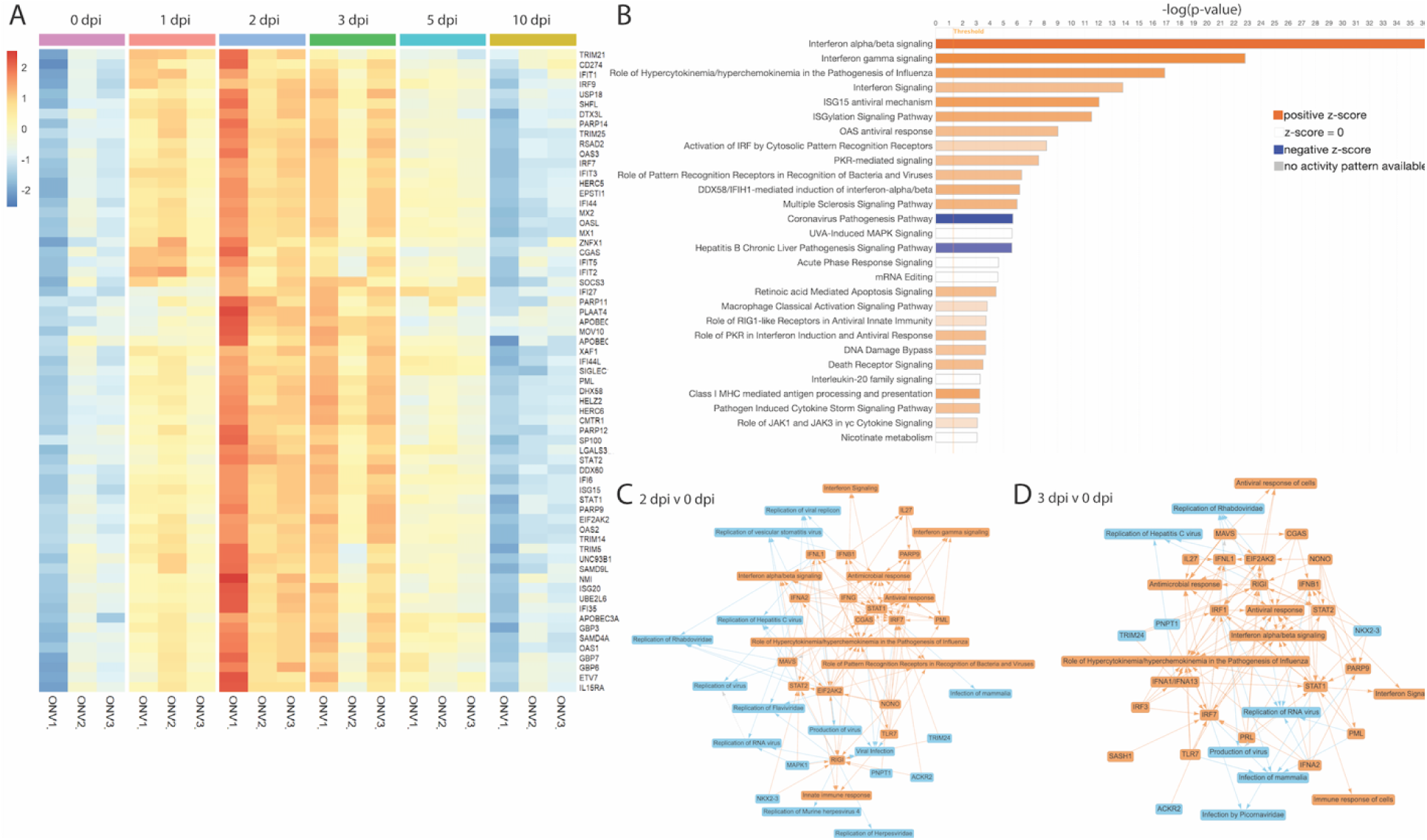
Transcriptional analysis of changes in PBMCs following ONNV infection. **(*A*),** Heat map of selected differentially expressed (DE) genes between 0 and 2 dpi (FDR p<0.05 and FC>2). **(*B*),** Pathway analysis of the top enriched pathways between 0 and 2 dpi (FDR p<0.05 and |FC|>2). Graphical summary of the top hits for pathways and transcripts that are altered between 0 and 2 dpi **(*C*)** and between 0 and 3 dpi **(*D*)** (FDR p<0.05 and |FC|>2) generated using Ingenuity Pathway Analysis software. Colors in all plots encode z-scores that are predicted upregulated in red/orange or predicted downregulated in blue.

Consistent with these upstream cues, we observed enrichment of pathways involving MHC class I antigen processing and presentation, neutrophil degranulation, phagosome formation, and macrophage classical activation signaling **(S Figure 6)**. These pathways point to the influx of neutrophils, monocytes and macrophages, and dendritic cells, and corroborate our WBC analysis **(S Figure 3)**. Although innate immune programs predominated at 2 dpi, early adaptive signatures were already detectable, including IFN-gamma responses, TRIM21 intracellular antibody signaling pathway, and Th1 polarization, suggesting initial activation of T and NK cells with emerging cytotoxic effector programming **(S Figure 6)**. The overall transcriptional landscape changes significantly by 10 dpi, largely returning to baseline for initial interferon responses. Top transcriptome DE genes that were linked to cellular recovery, recruitment of myeloid cells, and engulfment of cells **(S Figure 7A-C)**.

Our previous transcriptomics study in PBMC isolated from MAYV-infected rhesus macaques (26) revealed similar themes in innate immune signaling. Both infections triggered overlapping innate immune signatures at 2 dpi, including upregulation of MX1/2, OAS1/2/3, RIGI, ISG15, and SIGLEC1 **(Figure 6A)**, which normalized by 10 dpi. Despite some inter-animal variability, overall patterns were highly comparable between viruses. IPA identified interferon responses as top-enriched for both, while ONNV showed stronger activation of type I/II interferon pathways. In contrast, MAYV was characterized by additional regulation of EIF2 signaling (translation control) and enrichment of mTOR, IL-12, and B cell signaling pathways, reflecting subtle divergence in downstream immune programming between the two alphaviruses.

**Figure 6.**
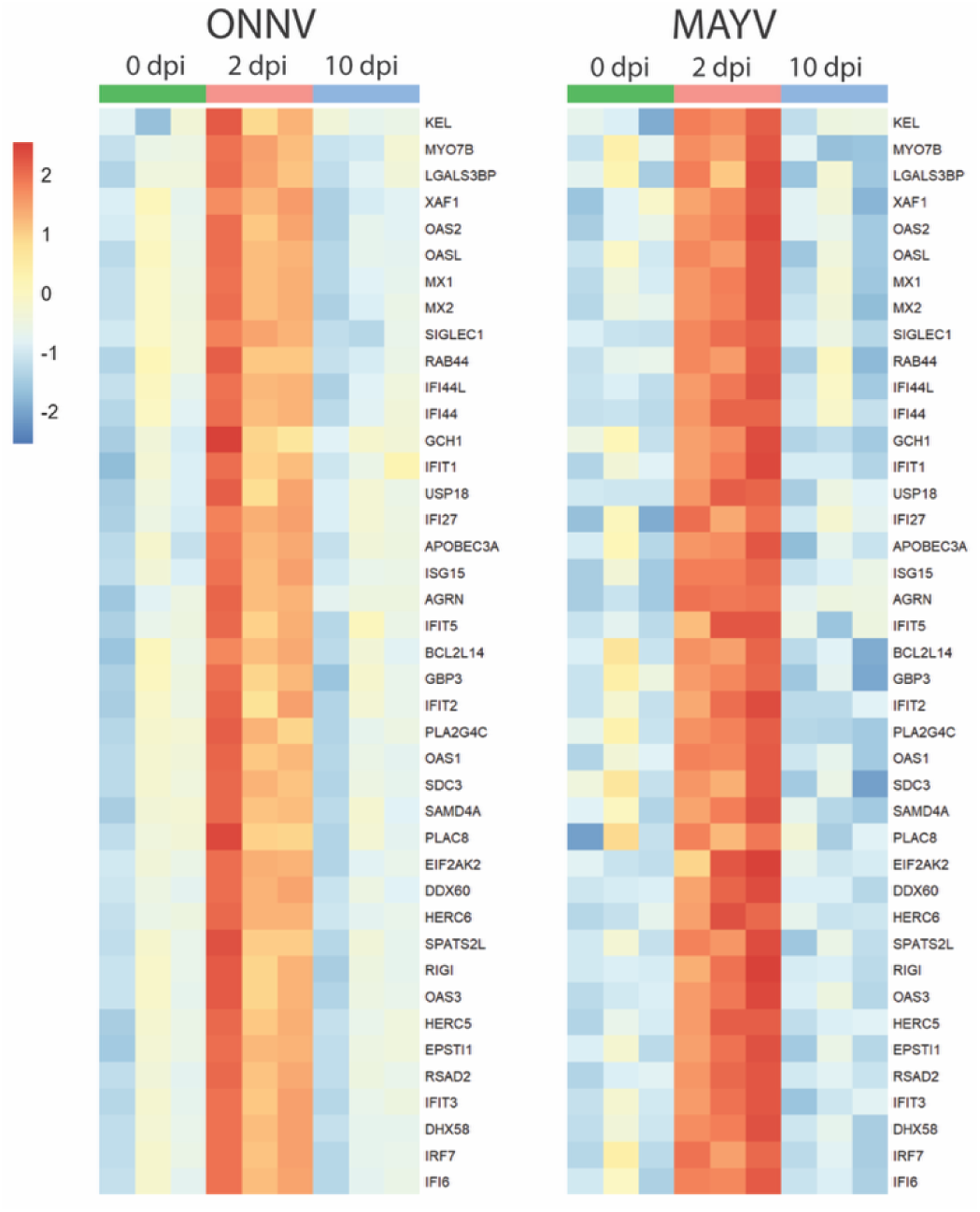
Transcriptional analysis of changes in PBMCs for shared genes between ONNV and MAYV rhesus macaque models. Previously published MAYV rhesus macaque (RM) RNA sequences were realigned to the latest RM genome that was used for this analysis (26). Heat map of top up-regulated genes between 0 and 2 dpi (FDR p<0.05 and FC>4) shared between ONNV and MAYV. Colors encode z-scores that are more upregulated in red/orange or more downregulated in blue.

### 4.5 Cellular innate immune signatures peak with ONNV viremia in rhesus macaques

Activation of monocytes, macrophages, and dendritic cells following infection of RM with many different arboviruses has consistently been shown to correlate with increased levels of CD169+ (SIGLEC1) staining by flow cytometry, and CD169 was a top-regulated DE gene (2 dpi: FC 3.988) identified in our transcriptomics analyses **(Figure 5B)** (26, 47). To understand the kinetics of these innate immune responses in our ONNV-infected RMs and to validate the DE analysis findings, we quantified the frequency of total and activated (CD169+) monocytes and dendritic cells in longitudinal PBMC samples **(Figure 7)**. After a repeated-measures ANOVA with the Geisser–Greenhouse correction, followed by Sidak’s multiple comparisons test was applied to the data, we found that the classical, intermediate, and nonclassical monocytes showed significant activation that coincided with viremia kinetics peaking between 2 and 5 dpi, and then returning to pre-infection baseline levels by 10 dpi **(Figure 7A–7C)**. Myeloid dendritic cells followed a similar pattern and level of significance; however, we did not detect activation of blood-derived plasmacytoid dendritic cells **(Figure 7D, E)**. While increases in activation for monocyte and myeloid dendritic cell subsets were observed, there were no significant changes in the total frequencies of any of these cell populations **(Figure 7A–7E)**.

**Figure 7.**
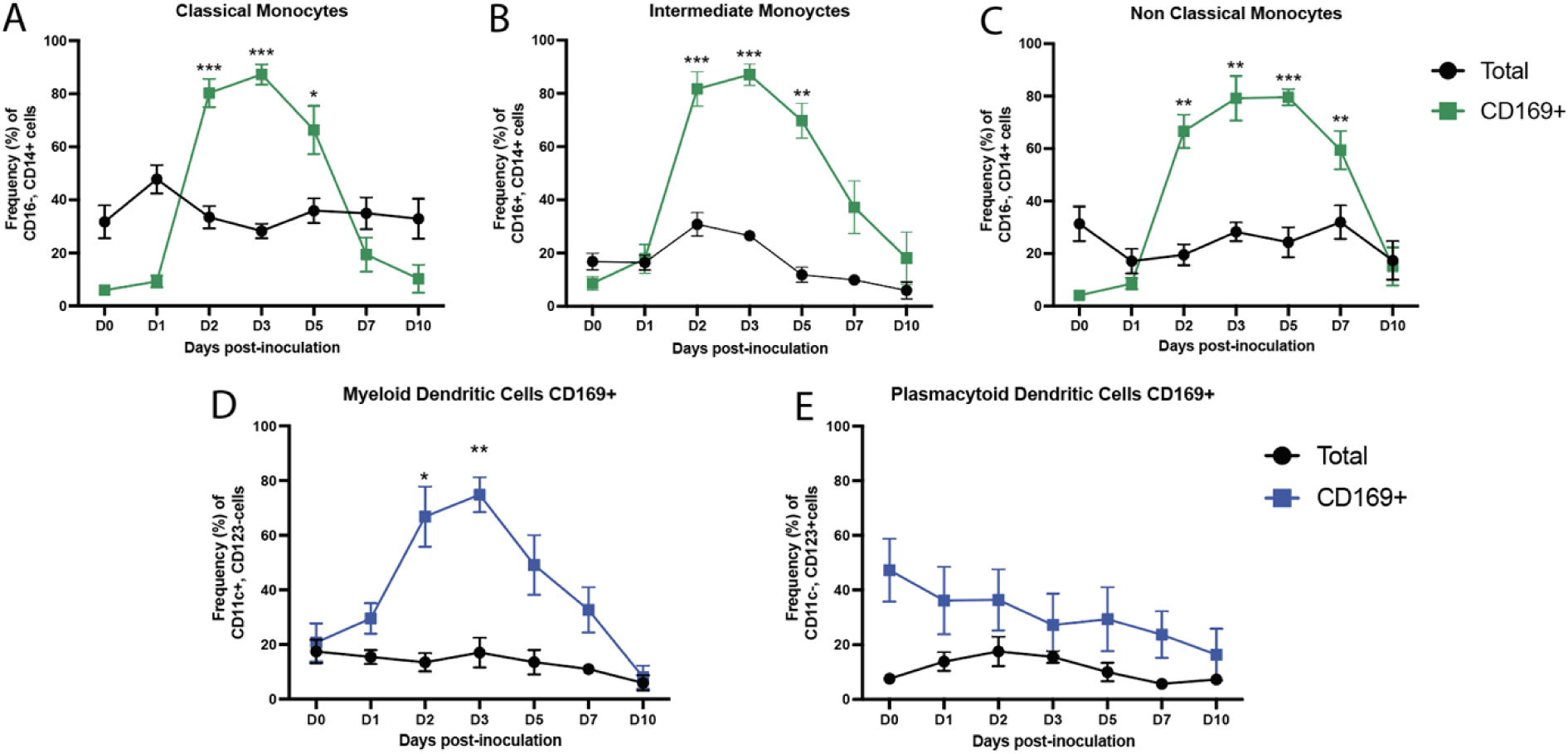
Longitudinal analysis of innate immune cell phenotypes and activation in ONNV-infected rhesus macaques. Whole blood was processed to remove plasma and lyse red blood cells prior to staining with antibodies directed against lineage and activation specific markers. Samples were analyzed by flow cytometry to assess innate immune cell subsets and activation status. Frequencies of total and activated (CD169⁺) classical monocytes ***(A)***, non-classical monocytes ***(B)***, and intermediate monocytes ***(C)*** were calculated. Dendritic cells (CD16⁻/CD14⁻) were further subdivided into myeloid dendritic cells ***(D)*** and plasmacytoid dendritic cells ***(E)***, with both total and activated (CD169⁺) populations measured. Lines represent mean frequencies of the six animals and error bars represent the standard error of the mean. Longitudinal changes were analyzed in Prism using a two-way repeated-measures ANOVA (time × treatment) with the Geisser–Greenhouse correction, followed by Sidak’s multiple comparisons test to compare each post-challenge day to baseline (day 0) within groups. Significance is shown as: p < 0.05 (*), p < 0.01 (**), p < 0.001 (***); all other comparisons were ns (p ≥ 0.05). Analysis was performed in GraphPad prism v10.2.0.

### 4.6 Proliferating T cell subsets were detected in response to ONNV infection in rhesus macaques

T cell responses are important for controlling alphavirus-mediated infection and disease (48, 49, 50), with CD4+ T cells in particular having also been shown in mice to mediate early infection-associated joint disease (50, 51). To characterize T cell responses following ONNV infection, we performed multiparametric flow cytometry on longitudinal rhesus macaque whole blood samples **(Figure 8A–E)** using a well-validated antibody panel (26, 47). Ki67⁺ central memory (CM) CD4⁺ T cells increased beginning at 3 dpi, with a steady rise between 7 and 10 dpi (**Figure 8A)**. One-way repeated-measures ANOVA showed a trend towards significance for the effect of time (p = 0.22). However, post-hoc Sidak comparisons vs baseline were not significant, likely due to high variability in the levels between animals. No increase in Ki67 staining was detected for either effector memory (EM) CD4+ T cells or naïve CD4+ T cell populations, which is similar to the responses observed in MAYV-infected NHP (26). Granzyme B expression by CD4⁺ EM T cells increased between 5–14 dpi, peaking at 7 dpi **(Figure 8B)**, although again, the overall ANOVA did not detect a significant time effect (p = 0.48).

**Figure 8.**
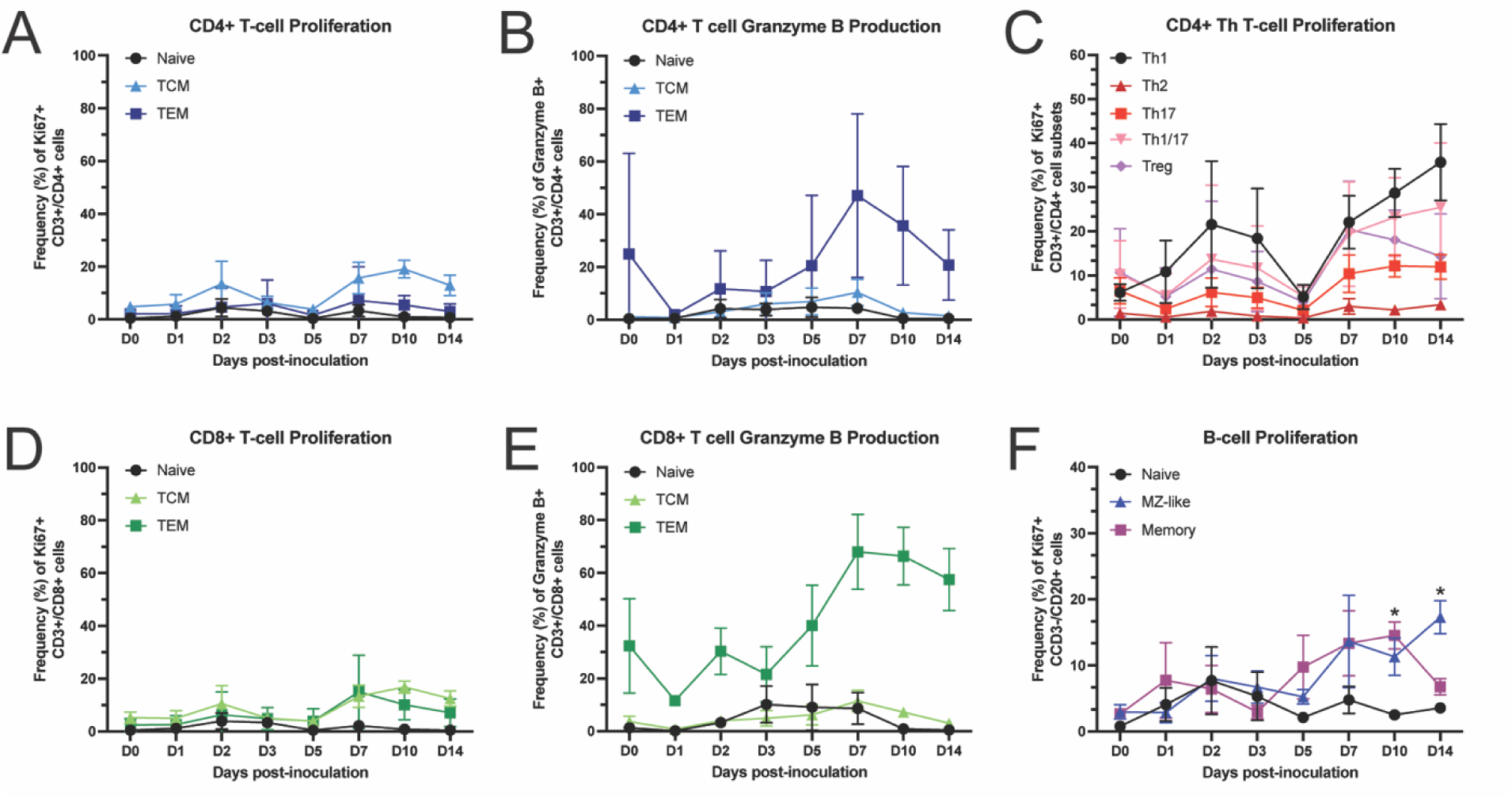
Kinetics of T cell proliferation and granzyme B expression in peripheral blood. Whole blood from ONNV+ rhesus macaques was processed to remove plasma and lyse red blood cells prior to staining with antibodies against lineage (CD3, CD4, CD8), proliferation (Ki67), and activation marker (granzyme B) followed by flow cytometry analysis. Changes in the longitudinal frequency of proliferating naïve, central memory, and effector memory CD4+ T-cells **(A)** and CD8+ T-cells **(D)** as well as granzyme B expression (granzyme B+) by CD4+ T **(B)** and CD8+ T cells **(E)** are shown and proliferating CD4⁺ T helper subsets (Th1, Th2, Th17, Th1/17) **(C)**. B cell (CD3-/CD20+) responses were separated into naïve, marginal zone (MZ)-like, and memory subsets and quantified for Ki67+ (proliferation). Lines represent mean frequencies of the six animals and error bars represent the standard error of the mean. Longitudinal changes were analyzed in Prism using a one-way repeated-measures ANOVA (time) with the Geisser–Greenhouse correction, followed by Sidak’s multiple comparisons test to compare each post-challenge day to baseline (day 0) within groups. Significance is shown as: p < 0.05 (*); all other comparisons were ns (p ≥ 0.05). Analysis was performed in GraphPad prism v10.2.0.

We further characterized T-helper subsets that proliferate following ONNV infection using CCR6 and CXCR3 to define Th1, Th2, Th17, and Th1/17 responses. The predominant subset that proliferated between 7-14 dpi was Th1 **(Figure 8C)**, which are known to produce interferon (IFN)-gamma, interleukin (IL)-2 and tumor necrosis factor (TNF)-beta, to evoke cell-mediated immunity and phagocyte-dependent inflammation (52, 53). Th1/17 cells (CCR6⁺CXCR3⁺) also expanded modestly, while proliferation of Th1/17 was minimal. Conversely, Th2 cells showed no detectable proliferative response. Regulatory T cells (CD25⁺CD127⁻) transiently increased in proliferation at 7 dpi but returned to baseline by 14 dpi **(Figure 8C)**.

CD8⁺ T cells, which are instrumental in promoting viral clearance and modulating disease, also displayed dynamic changes following ONNV challenge **(Figure 8D, E)**. Both CM and EM CD8⁺ T cell subsets showed modest increases in Ki67 expression around 7 dpi, plateauing by 14 dpi. In contrast, granzyme B expression in EM CD8⁺ T cells expanded more robustly, with frequencies steadily rising to reach peak values at 14 dpi. Although statistical analysis did not identify significant differences versus baseline, the temporal pattern suggests gradual expansion of activated cytotoxic CD8⁺ T cells during acute infection.

In addition to T cell responses, we evaluated B cell activation across naïve, marginal zone (MZ)-like, and memory compartments **(Figure 8F)**. Unlike T cells, clear differences emerged: memory B cells significantly increased at 7 dpi compared to baseline (p = 0.02), and MZ-like B cells significantly expanded at 10 dpi vs day 0 (p = 0.016). Naïve B cell frequencies remained stable throughout the infection time course with a slight elevation at 10 and 14 dpi. These findings demonstrate that ONNV infection elicits coordinated cellular immunity in rhesus macaques, characterized by proliferative Th1 responses, activation of cytotoxic CD8⁺ T cells, and proliferation of both memory and MZ-like B cell compartments, although high inter-animal variability tempered the statistical resolution of several T cell subsets.

### 4.7 Virus-specific antibodies emerge by 5 dpi and broaden in neutralization and effector functions by 14 dpi

To help identify the correlates of protection, we defined the kinetics, magnitude, and breadth of antibody development following infection with ONNV_0804_. Consistent with CHIKV and MAYV infection dynamics, we detected ONNV-specific IgM as well as IgG antibodies as early as 5 dpi in all six macaques **(Figure 9A)**. IgM antibody responses were more robust initially as expected during the acute infection period; however, the IgG responses are likely to surpass the IgM levels at later timepoints as seen in other arboviral infections **(Figure 9A)**. ONNV-neutralizing antibodies were detected as early as 5 dpi in all six macaques using plaque reduction neutralization assays **(Figure 9B)**. These neutralizing antibody levels reached 50% plaque reduction neutralization titers (PRNT50) of 3.5×10^3^ - 4.6×10^4^ by 14 dpi **(Figure 9B)**.

**Figure 9.**
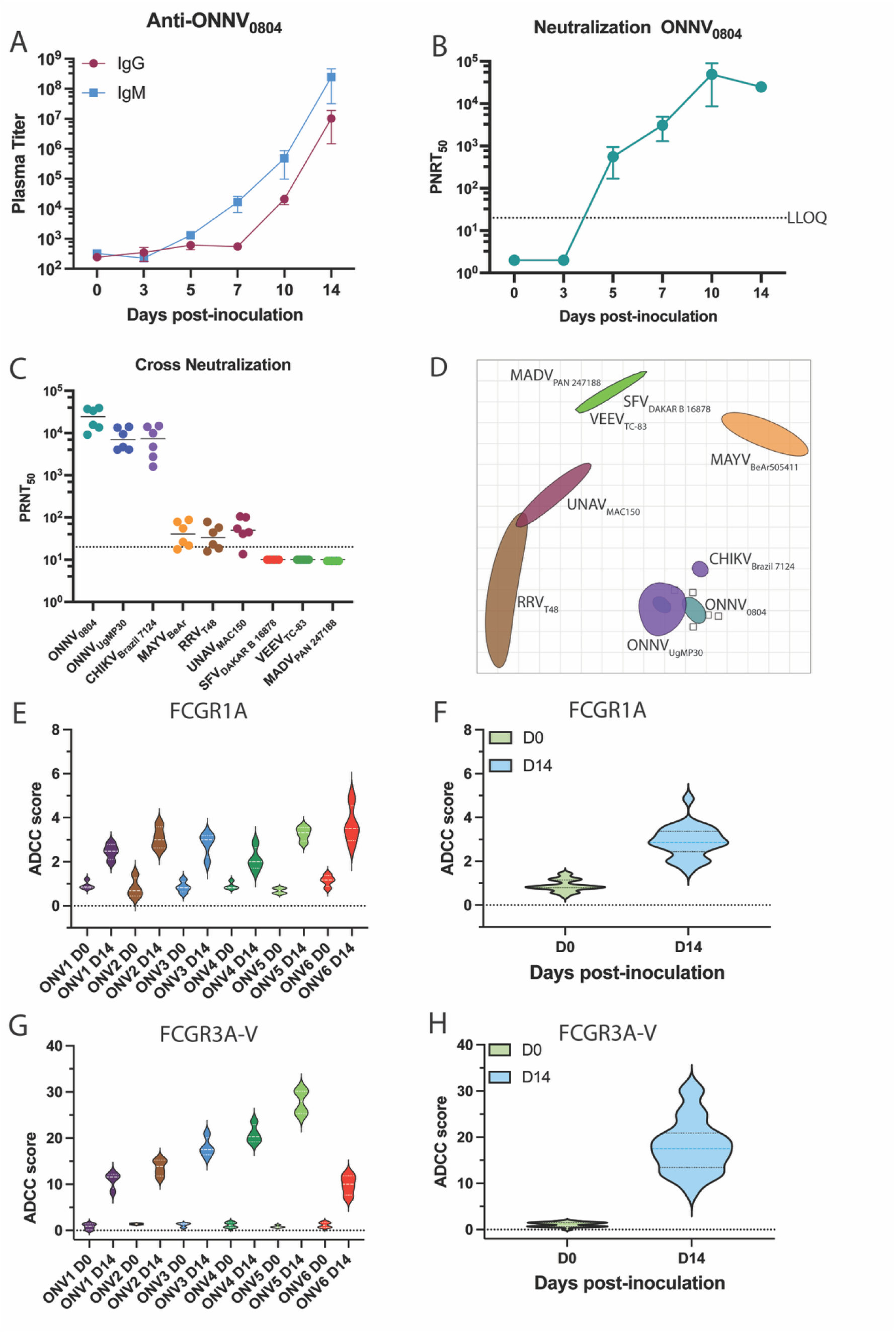
Characterization of ONNV-specific antibodies and analysis of cross-reactive breadth. ***A,*** Development of ONNV-binding, IgM and IgG isotype antibody titers were quantified in macaque plasma at 0, 5, 7, and 10 dpi by ONNV antigen specific ELISA. The LOD was a 1:50 plasma dilution with undetectable values graphed as half the LOD. ***B,*** ONNV-neutralizing antibodies was quantified in plaque reduction neutralization assays using heat-inactivated macaque plasma at 0–5, 7, 10, and 14 dpi. 50% plaque reduction neutralization titers (PRNT_50_) were determined in non-linear regression on GraphPad prism. The LOD was a 1:20 plasma dilution and undetectable values were graphed as half of the LOD. Error bars are *SEM. **C,*** Breadth of antibodies of heat-inactivated macaque plasma that neutralized other relevant alphaviruses following ONNV infection (14 dpi) were characterized in cross-neutralization assays against CHIKV, MAYV, UNAV, RRV, VEEV, SFV, and MADV using day 0 plasma as a baseline control. The LOD was a 1:20 plasma dilution and undetectable values were graphed as half of the LOD. ***D,*** Antigenic cartography mapping of the antigenic distances between viruses based upon cross-neutralization data. ***E,F,*** Activation of FcγRIA signaling based on NFAT reporter assay separated by individual animals **(E)** and averaged across all animals **(F)** for D0 and D14. ***G,H,*** Activation of FcγR3A-V signaling for individual animals **(G)** and averaged across all animals **(H)** for D0 and D14. Violin plots represent data from two replicate experiments.

Finally, the antiviral breadth of neutralizing antibodies was determined using limiting dilution plaque neutralization assays against additional Semliki Forest antigenic complex viruses, including CHIKV, UNAV, MAYV, RRV, and Semliki Forest virus (SFV), and against the serologically unrelated alphaviruses VEEV and Madariaga virus (MADV) **(Figure 9C)**. Pre-infection plasma samples were found to be devoid of any neutralizing activity (PRNT50 < 20) against any of the viruses tested. At 14 dpi, cross-neutralizing antibodies were detected strongly against CHIKV, weakly against UNAV, MAYV and RRV, but not against SFV, VEEV, or MADV **(Figure 9C)**. Cross-neutralization at 14 dpi was greatest against viruses more antigenically related to ONNV, which is visualized using antigenic cartography **(Figure 9D)**. RM plasma clustered around ONNV strains due to highest neutralization potency with CHIKV positioned the closest to ONNV. MAYV, UNAV, RRV were positioned further away and, as expected, SFV, VEEV, and MADV are farthest away due to no detectable neutralization against these viruses **(Figure 9D)**. In summary, our findings indicate that ONNV-specific antibodies develop as early as 5 dpi and expand in both magnitude and breadth, with the capability to neutralize only related SFV complex alphaviruses.

To further characterize the functional antibody landscape following ONNV challenge, we examined Fcγ-receptor (FcγR) activation profiles for plasma collected at 0 and 14 dpi. FcγR interactions provide comprehensive immune responses by modulating adaptive immune responses, activation of T- and B-cells, facilitating direct elimination of virus-infected cells, as well as enhancing long-term immunity. While neutralizing antibodies remain the classical correlate of protection for most alphaviruses, many monoclonal antibodies achieve maximal potency only when FcγR-dependent effector functions are engaged (54, 55, 56, 57). For this analysis, we focused on three receptors that represent distinct effector pathways: FCGR1A (CD64), a high-affinity receptor expressed primarily on monocytes and macrophages that mediates antibody-dependent phagocytosis and antigen uptake, FCGR3A-V158 (CD16a), a low-affinity activating receptor predominantly found on natural killer (NK) cells that drives antibody-dependent cellular cytotoxicity (ADCC), and finally FCG2A-H an intermediate-affinity receptor expressed on myeloid cells and platelets that contributes to antibody-dependent phagocytosis and inflammatory signaling. All 6 animals showed >2-fold ADCC score at D14 for both FCGR1A and FCGR3A-V, but no evidence of activation for FCGR2A-H **(Figure 9 E, F, G, H)**. Notably, ADCC scores for FCGR1A and FCGR3A were moderately correlated, with ONV5 exhibiting the highest responses for both receptors, consistent with this animal’s overall viral kinetics and clinical profile. Overall, these data demonstrate that ONNV infection generates a rapid, cross-reactive, and functionally competent antibody response, providing a foundation for understanding humoral correlates of protection and vaccine design.

## 5. Discussion

To establish a translational O’nyong-nyong virus (ONNV) model, we previously constructed an infectious clone for a contemporary strain, ONNV_0804_ that was isolated and sequenced from a febrile patient. We evaluated strain-specific differences in mice using this new infectious clone and a laboratory strain UgMP30 and determined that infection with ONNV_0804_ caused earlier and more severe disease and higher viremia (24). Building on these results, herein we infected six rhesus macaques (RM) with 10⁵ PFU ONNV_0804_ to assess pathogenesis and immune dynamics in a primate system. Our findings from this study demonstrate that ONNV elicits robust innate and adaptive immune responses in rhesus macaques, revealing both shared and distinct features compared to other clinically important arthritogenic alphaviruses such as CHIKV and MAYV.

In this study, all six animals had productive infection dynamics with peak viremia observed at 2 dpi, closely mirroring CHIKV and MAYV infection kinetics in RM (26, 30, 36). Clinically, ONNV infection in humans presents with fever, rash, malaise, myalgia and arthralgia within the first week of infection (1, 58). The ONNV-infected RM developed a macular to maculopapular rash and dorsal wrist edema that developed as early as 7 dpi. Histology on these rashes displayed a range of perivascular lymphocytic aggregates to lymphoplasmacytic dermatitis with hyperkeratosis and dermal edema. Although fever could not be accurately measured due to study design, this sign is rarely detected in RM models of arboviral infection (26, 47, 59). Three of the rhesus macaques developed lymphadenopathy, which is another clinical feature of ONNV infection (58).

In mice, ONNV infection has been shown to disseminate to various tissues, including dermal fibroblasts, muscles, heart, and brain, and viral RNA can persist at least up to 43 dpi (24). In rhesus macaques, ONNV similarly disseminated broadly, infecting skeletal muscle, joints, peripheral nerves, lymphoid tissue, skin, multiple brain lobes, and primary organs. Since infectious MAYV was detected in select tissues at 10 dpi (26), we attempted a similar approach but, unfortunately, infectious virus isolation from key tissues of interest (joint, muscle, CNS), was unsuccessful, which we attribute to the later study endpoint (14 dpi).

Histologic analysis revealed that wrists and fingers were the most affected joints in the ONNV infected animals, with lymphoid infiltrates primarily localized to periarticular adipose tissue and loose connective tissue, including adipose and collagenous connective tissue. These lesions were characterized by mononuclear and lymphoplasmacytic inflammation without overt synovial destruction. The distribution of inflammation suggests early periarticular involvement rather than established synovitis, consistent with an acute or resolving phase of infection. In human biopsies from other arthritogenic alphavirus infections, vascular proliferation, synovial hyperplasia, and mononuclear perivascular aggregates to inflammation extending into periarticular tissues have been reported, chiefly consisting of monocyte lineage cells with or without CD4+ T helper cells (60, 61). Similarly, in mouse models, synoviocytes appear to be particularly susceptible to ONNV infection with histologic correlates of notable muscle degradation, synovitis, and tenosynovitis surrounding the tendon sheath (24, 62, 63). Compared with these models, the rhesus macaque lesions observed here were more limited in scope, potentially reflecting the early timepoint examined, differences in sampling sites, or species-specific variation in ONNV pathogenesis. While our study was limited to six animals and a relatively short infection window, the consistent periarticular inflammation and detection of viral RNA within joint tissues highlight early immunopathologic processes that may precede chronic arthritic disease. Although our immune analyses were limited to peripheral blood, the timing of histologic inflammation suggests that localized immune activation within joint tissues could contribute to early pathogenesis. Future work should focus on later timepoints to determine whether these early lesions evolve into chronic arthritis, as well as on spatial analyses to identify specific inflammatory and stromal cell types driving local pathology.

Having established clinical and pathological correlates, we next examined innate immune activation. Alphavirus infection typically induces potent type I interferon responses essential for viral clearance. Mice lacking type I IFN signaling exhibit high mortality, with type II IFN responses appearing to play no role in mice’s survival following ONNV infection (24, 25). Indeed, in RM transcriptomic analysis identified a robust type I interferon response consistent with classical antiviral signaling with secondary type II IFN activation. Upregulation of transcripts like SIGLEC-1 (CD169+) was confirmed via flow cytometry, which peaked with kinetics similar to viremia. These signatures, together with enrichment of antigen presentation and macrophage activation pathways, underscore the rapid engagement of innate antiviral defenses that likely constrain systemic replication, paralleling early responses described in CHIKV and MAYV infection.

The role of adaptive immunity in CHIKV and MAYV infection is well characterized with B cells dominating the late-acute phase adaptive immune responses and contributing to viral clearance (48, 64). Antibodies are typically against the E-proteins and provide neutralizing and cross-neutralizing protection. Similar to MAYV RM models, we showed longitudinal antibody responses and that the breadth for cross reactivity was more limited when compared to MAYV. ONNV-elicited antibodies engaged Fc gamma receptors (FcγRs), indicating a role for Fc-dependent effector mechanisms in protection. While in other systems, such as SARS-CoV-2 and dengue virus, FcγR engagement has been associated with either protection or enhancement, respectively, our results indicate beneficial Fc-mediated activation, particularly through FCGR3A-V and, to a lesser degree, FCGR1A (65). These findings underscore the importance of antibody quality, not just quantity, in defining correlates of protection, and this information could be useful in guiding vaccine design.

T cells have been shown to have more damaging effects during chronic joint infection in CHIKV, with Th1 polarized CD4+ T-cells as a driver of both acute and chronic arthritic joint pathology (66, 67). In ONNV-infected mice, CD4+ T-cells, CD8+ T-cells, and monocytes were present within joint tissues during peak foot-pad swelling, and specifically, CD4 depletion led to decreased joint pathology (63). Although our analysis focused primarily on proliferative responses, we observed Th1 polarization of CD4⁺ T cells, consistent with transcriptomic enrichment of IFN-γ signaling and Th1 pathways. Given the concurrent IFN-γ–driven transcriptomic signature and Th1 polarization, these findings suggest a coordinated antiviral response that may also contribute to immunopathology during persistent joint inflammation. However, future studies are needed to investigate the cellular-specific immune infiltration at extended timepoints.

## 6. Conclusion

Recent advances in anti-alphavirus immunotherapeutics, vaccines, and small molecule antivirals coupled with the increasing geographic overlap of alphaviruses have highlighted the need for testing the efficacy breadth across alphavirus species. Thus, there is a critical need for the development of pre-clinical platforms for testing alphavirus vaccines and antivirals due to the increased global threat of alphavirus emergence. The current study establishes a nonhuman primate model of O’nyong-nyong virus infection, recapitulating many of the hallmark features of human disease, such as viremia and tissue distribution, evidence of early arthritogenic disease, lymphadenopathy, and skin rashes in most animals. Although the absence of longitudinal studies in humans limits our understanding of the long-term impact of ONNV infection, chronic symptoms of arthritogenic alphaviruses are known to greatly impact quality of life and productivity due to long-term incapacitating polyarthralgia. Our results have important implications for studying therapeutics to mitigate this disease and additionally extend not only to vaccines that provide protection against ONNV but also provide cross-protection to closely related alphaviruses such as the recent FDA-approved CHIKV vaccine, VIMKUNYA. By establishing a nonhuman primate model of O’nyong-nyong virus infection, our study bridges a critical gap in understanding the pathogenesis and immune responses to this historically neglected but reemerging alphavirus.

## 7. Methods

### 7.1 Animal ethics statement

All experimental protocols were approved by the Institutional Animal Care and Use Committee (IACUC) of the Oregon National Primate Research Center (ONPRC), which is accredited by the Assessment and Accreditation of Laboratory Animal Care (AAALAC) International. All procedures in this study complied with the ethical standards outlined in the Animal Welfare Act, as enforced by the United States Department of Agriculture, and followed national guidelines for the care and use of laboratory animals. The study also adheres to the ARRIVE guidelines for animal research (68).

### 7.2 O’nyong-nyong 0804 (ONNV_0804_) infectious clone construction

The O’nyong-nyong 0804 infectious clone was constructed, as previously described, using the sequenced genome from a 2017 clinical isolate of a febrile patient from Uganda (24). The entire genome was synthesized as approximately seven fragments, each ∼1.7kb in length, and containing 20bp of overlapping homology for NEBuilder assembly. The seven fragments, plus the infectious clone vector with overlapping homology, were subjected to NEBuilder assembly as per NEB’s protocol using equal molar concentrations of all fragments. Constructs were validated by Oxford nanopore sequencing (Eurofins).

### 7.3 Cell culture and Virus preparation

Vero cells (ATCC CCL-81) were propagated at 37°C and 5% CO2 in Dulbecco’s Modified Eagle Medium (DMEM; Thermo Scientific) containing 5-10% fetal calf serum (FCS; Thermo Scientific) supplemented with 1X penicillin-streptomycin-glutamine (PSG; Life Technologies). Ae*des albopictus* C6/36 cells (ATCC CRL1660) were grown at 28°C with 5% CO2 in DMEM containing 10% FCS and 1X PSG. Alphaviruses Mayaro (MAYV_BeAr505411,_ NR-49910), Una (UNAV_MAC150_, NR-49912), ONNV_UgMP30_ (NR-51661), Ross River (RRV_T-48_, NR-51457), and VEEV_TC-83_ (NR-63) were obtained through BEI Resources, NIAID, NIH. Semiliki Forest virus (SFV_Dakar B 16878_) and Madariaga virus (MADV_PAN 243178_) were obtained from the World Reference Center for Emerging Viruses and Arboviruses at The University of Texas Medical Branch-Galveston. Viruses were prepared as previously described (24, 26). Chikungunya virus (CHIKV_181/25_) was generated from an infectious clone as previously described (69, 70). Viruses were propagated as described previously, and titered on Vero cells using limiting dilution plaque assays in 48-well plates (24). Infected cells were incubated for 2 hours under continuous rocking at 37°C with 5% CO₂, then overlaid with a 2:1 mixture of DMEM-5 containing 0.3% high/low viscosity carboxymethyl cellulose (CMC-DMEM) (Sigma). Cells were fixed with 3.7% formaldehyde and stained with 0.2% methylene blue at 48 hpi for MAYV_BeAr505411_, UNAV_MAC150_, VEEV_TC-83_, CHIKV_181/25_ or at 72 hpi for ONNV_UgMP30_ and ONNV_0804_. Plaques were used to calculate viral titers in plaque-forming units (PFU) per mL. Virus stocks for all experiments were passaged 1 or 2 times to limit acquired mutations and were sequence-validated as described below (24).

### 7.4. Nonhuman primate experiments

Six adult rhesus macaques (*Macaca mulatta*) three males ages 4, 4 and 5 years and three females ages 8, 11, and 12) were included in this study. This is consistent with ethical, logistical, and cost constraints typical of nonhuman primate infectious disease model development (26, 43, 47). Although budget and availability constraints make experiments with NHP underpowered compared to mouse studies, our prior experiments have shown statistically significant effects of Ab treatment on CHIKV-infected rhesus macaques (RM) (40) using cohort sizes of 4 or 5. Assuming similar results in the proposed studies, cohorts of at least 4 NHP will give us 80% power to detect a 1.3 log10 change in viral RNA (a=0.05), and observe significant effects in pathology (based on severity of inflammatory infiltrate and area affected). Thus, our cohort size of n=6 allowed for the generation of robust, high-quality data. After data collection, a post-hoc analysis was performed using the observed variability in the primary endpoint, peak plasma viremia (log₁₀ copies/mL) analyzed with R Studio. The mean ± SD peak viremia was 5.99 ± 0.88 log₁₀ copies/mL, with a 95% confidence interval of 5.07–6.91. Under these variance assumptions, a two-sided α=0.05 test with n=6 animals provides ∼80% power to detect differences of ≥1.57 log₁₀ copies/mL between groups.

All NHPs were healthy, average-weight animals that were fed a standard chow diet (Purina Mills) and had access to water *ad libitum*. Animals were inoculated subcutaneously with a total of 10^5^ plaque-forming units (pfu) of ONNV_0804_ diluted in 1 mL of physiologic saline as determined by a mouse model of ONNV (24). The inoculum was distributed as ten total 100 µl injections at the dorsal (back) hand in both radial and medial positions, the radial and medial aspects of the dorsal wrist, and one injection near the proximal antebrachium (forearm) of each arm. This anatomical distribution was selected to mimic mosquito bites and promote uniform subcutaneous absorption and systemic dissemination of the virus, while ensuring consistency. Post-challenge animals were monitored daily for clinical signs of disease, including temperature, body weight changes, rash, edema, and signs of discomfort. Animals were placed in single housing following inoculation to control for viral shedding and potential reinfection to co-housed animals. Enrichment was provided, and each animal had continuous full visual contact with the other animals. Blood was collected at indicated timepoints (0, 1–5, 7, and 10 dpi) for monitoring by complete blood count, serologic studies, cellular activation (PBMCs) assays, and transcriptional changes following challenge. Rhesus macaques were humanely euthanized at 14 dpi and complete necropsies were performed. Three representative tissue sections (∼1cm^3^) from joint, muscle, lymphoid, major organs, nervous system, and reproductive tissue were collected into 1mL TRIzol reagent (Invitrogen) for RNA isolation, preserved in RNAlater, or fixed in 4% paraformaldehyde for histopathology.

### 7.5. Enzyme-linked immunosorbent assay (ELISA)

Enzyme-linked immunosorbent assays (ELISAs) were conducted on high-binding 96-well plates (Corning) coated for 3 days at 4°C with 1×10⁶ plaque-forming units (pfu) per well of purified ONNV_0804_ in PBS. Plates were then blocked with 5% milk in PBS containing 0.05% Tween-20 before incubating with serial five-fold dilutions of heat-inactivated plasma, starting at a 1:50 dilution. Detection was performed using horseradish peroxidase (HRP)-conjugated anti-monkey IgG or IgM secondary antibodies (Rockland) at 1:1000, followed by OPD substrate (0.05 M citrate buffer, 0.4 mg/mL o-phenylenediamine, 0.01% hydrogen peroxide, pH 5; Life Technologies). After 7 minutes, the reactions were stopped with 1 M HCl. Absorbance was measured at 490 nm using a Synergy HTX Microplate Reader (BioTek, Winooski, VT). Endpoint titers were calculated using a Log/Log transformation method, and results were analyzed and graphed in GraphPad Prism v10.2 software.

### 7.6. Plaque reduction neutralization (PRNT) assays

Plaque Reduction Neutralization Tests (PRNTs) were conducted as previously described to assess plasma antibody neutralizing activity. Plasma samples were heat-inactivated at 56°C for 30 minutes, and then serially diluted five-fold in 2% FBS/DMEM, starting at a 1:20 dilution. Diluted plasma was incubated with 10^7^ PFU per well of ONNV_0804_ at 37°C for 2 hours. The virus–plasma mixtures were then transferred onto confluent monolayers of Vero cells in 12-well plates and incubated for 2 hours with intermittent rocking. After infection, cells were overlaid with carboxymethylcellulose (CMC) in 5% FBS/DMEM. Viruses were incubated, fixed, and stained as described above. The 50% plaque neutralization titers (PRNT50) were calculated by non-linear regression analysis using GraphPad Prism v10.2 software after determining the percent of plaques at each dilution relative to control wells containing no plasma.

### 7.7. Viral RNA detection

Blood, urine, and cerebrospinal fluid (CSF) RNA was isolated using Promega Maxwell prep using the Total Nucleic Acid kit according to manufacturer’s protocol. Tissue sample RNA was isolated using Zymo Research Direct-zol RNA Miniprep Kit (Cat. R2050) according to the manufacturer’s protocol. Tissue sections were collected into tubes containing 1 mL of TRIzol, and approximately 250 μL of silica beads (VWR 48300–437). The samples were homogenized using a Precellys 24 homogenizer (Bertin Technologies), pulsing for two cycles of 45 seconds on and 30 seconds off. ONNV_0804_ RNA levels were measured by a one-step quantitative real-time reverse transcription polymerase chain reaction assay (qRT-PCR) using TaqMan One-Step RT-PCR Master Mix (Applied Biosystems) as described previously (24). Primer/probe sequences for the detection of ONNV_0804_ include Forward: 5’-CCCACAGCATGGCAAAGAAC; Reverse: 5’CTGGCGGCATATGCACTTCT; and TaqMan probe: 5’ Fam-ACGTACGTCCATACCACAG-MGB. For RNA standards, RNA was isolated from a purified, tittered stock of ONNV_0804_. Final numbers are reported as a log transformation.

### 7.8. Histopathological analysis

Representative tissue sections were collected, fixed in 4% paraformaldehyde for 24 hours, moved to 70% ethanol at 4°C, embedded in paraffin, and 5 μm sections were prepared and stained with H&E on a Leica ST5020 Autostainer. All slides were assessed for histopathology on Leica DFV495 or Leica DM3000 light microscopes by M.D. /Ph.D immunopathologist (T.F.A.) and board-certified veterinary pathologists (C.S.L. and M.K.A.). Sections from musculoskeletal tissues were additionally evaluated for inflammation based upon a histologic grading rubric (**S Table 2, S Figure 5**), quantifying perivascular lymphoid aggregates and inflammation with note of tissue types affected and any synovial hyperplasia or hypertrophy. As expanded upon in S Table 2, severity scores indicated the absence of any changes (0), scarce (1-2), minimal (3-4), mild (5-6), or severe (9-10) disease. All slides were scanned using the Zeiss Axio scanner at 40x magnification, and figures were created using HALO (Indica Labs) v10.4.

### 7.9. Transcriptomics

Transcriptomic analysis was done as described previously (26). Briefly, Total RNA from rhesus macaque PBMC isolated using the TRIzol extraction method described above was prepared for transcriptomic analysis using the Illumina TruSeq Stranded mRNA Library Prep Kit (RS-122-2101, Illumina) as previously described (71). The library was validated using an Agilent DNA 1000 kit on a bioanalyzer. Samples were sequenced by the OHSU Massively Parallel Sequencing Shared Resource using an Illumina NovaSeq. Differential expression analysis was performed by the ONPRC Bioinformatics & Biostatistics Core as described previously (26). Pathway analysis was performed with the Integrated Pathway Analysis software package (QIAGEN Inc., https://www.qiagenbioinformatics.com/products/ingenuity-pathway-analysis), using a stringent cutoff for significant molecules of FDRp < 0.05 and |FC| > 1.5. The background reference set used was the dataset of all genes in the differential analysis.

### 7.10. Phenotypic analysis of peripheral blood mononuclear cells

Flow cytometry was performed to phenotype peripheral blood mononuclear cells (PBMCs) from whole blood samples as previously described, with a few modifications (47). Whole blood was centrifuged at 750 G for 15 minutes after which plasma was removed and replaced with an equal amount of PBS. The sample was then vortexed and aliquoted into three 100 μL samples for subsequent phenotype staining. The remaining sample was layered over lymphocyte separation medium (Corning) and centrifuged for 40 minutes at 2,000 rpm. Isolated peripheral blood mononuclear cells (PBMC) were washed in 1X PBS and frozen in liquid nitrogen at ∼2 million cells/mL in freezing medium containing 10%DMSO/40%FBS/50%RPMI.

For immunophenotyping, 100 μL of whole blood was incubated with 1 mL of ACK red blood cell lysis buffer (ThermoFisher) for 10 minutes. Cells were pelleted by centrifugation at 2,000 rpm and washed twice with 1X PBS to neutralize the lysis buffer. Fluorophore-conjugated antibodies targeting CD45, CD3, CD8a, CD14, CD16, CD11c, HLA-DR, CD56, CD123, and CD169 (activation marker) were used to stain for monocyte, macrophage, and dendritic cell populations. Monocytes/macrophages were identified as delineated as described previously (47). For T-cell characterization, staining included CD45, CD3, CD4, CD8a, CD25, CD28, CD95, CD127, CCR6, CXCR3, and intracellular Ki67 and Granzyme B. Ki67+ and Granzyme B+ populations were gated relative to each animal’s baseline (day 0). For both panels, CD45+ events were first gated to remove residual RBCs, followed by singlet gating prior to subset identification. Data were acquired on a BD Symphony A5 (BD Pharmingen) and analyzed using FlowJo v10.10.0.

### 7.11. Antigenic Cartography

Antigenic cartography was performed to observe the serologic relationships for ONNV-infected rhesus macaques against a panel of Alphaviruses using the 50% plaque reduction neutralization titers obtained as described above. Maps were constructed using the Racmacs package (https://acorg.github.io/Racmacs/, version 1.2.9, accessed on 16 October 2025) in R Studio (version 2024.12.1+563). Raw neutralization titers of each serum against each alphavirus in the panel were tabulated, and serum neutralization titers under the limit of detection were given a PRNT50 of 10. The map was computed for 500 optimizations with two dimensions, and the minimum column basis was set to none. The Bayesian method was used to perform 1000 bootstrap repeats with 100 optimizations per repeat of the tabulated neutralization titers. Finally, blobs were added to the map using “ks” algorithm method to represent the confidence levels of the neutralized viruses with the confidence level set at 0.68 and grid spacing set at 0.1. Each box of the grid is representative of a 2-fold difference in serum dilution.

### 7.12 Jurkat/Fc Receptor ADCC Reporter assay

To prepare antigen-presenting cells, 3,000 Huh-7.5 cells were plated in a 384-well plate (Corning 3764) in 20 μL DMEM supplemented with 10% fetal bovine serum, 1% penicillin-streptomycin, and 1% L-glutamine. The next day, cells were infected with ONNV (MOI 0.1). At 24 h post-infection, virus was inactivated using a UV Stratalinker 2400 (Stratagene). Lids were removed from the plates, and the machine was set to deliver 250 mJ/cm^2^, followed by 750 mJ/cm^2^. Growth medium was then robotically aspirated from assay plates such that 10 μL media remained per well. An 8-point dilution series of rhesus macaque plasma samples was robotically prepared by initially diluting plasma samples 1:10 in 2% (w/v) BSA in PBS, followed by 4-fold dilutions between concentrations in technical duplicate. 5 μL of the 8-point dilution series of rhesus macaque plasma was robotically transferred to assay plates, yielding a final plasma dilution range of 1:50 to 1:819,200. Dilution series of the control mAbs rEEV-179 and rEEV-179 with a LALA mutation, ranging from 10 μg/mL to 0.0006 μg/mL, were included on each assay plate. Nine thousand Jurkat/NFAT-Luciferase reporter cells, expressing either FCGR1A, or FCGR2A-H131, or FCGR3A-V158 were added to assay plates in 10 μL RPMI, supplemented with 10% ultra-low-IgG fetal bovine serum, 1% penicillin-streptomycin, and 1% L-glutamine. Assay plates were incubated at 37⁰C/5% CO2 for 24 h. Assay plates were removed from the incubator for 15 min to equilibrate to room temperature prior to adding 20 μL Brite-lite Luciferase Reagent (revvity). Luminescence was measured on an EnVision Xcite Multilabel Plate Reader (revvity), using the ultrasensitive luminescence measurement technology. Assay well data was normalized to aggregated negative control wells and expressed as antibody-dependent signal-to-background (S:B = sample well/NegC Avg). The ADCC score was defined as the S:B at a 1:3200 dilution of the plasma.

## Author Contributions

All authors have read and agreed to the published version of the manuscript.

**Hannah K. Jaeger** Conceptualization, Data curation, Formal analysis, Investigation, Methodology, Validation, Visualization, Writing – original draft.

**Michael Denton** Data curation, Formal analysis, Investigation, Methodology, Validation, Visualization, Writing – review and editing.

**Takeshi F. Andoh** Data curation, Formal analysis, Investigation, Methodology, Validation, Visualization, Writing – review and editing.

**Craig N. Kreklywich** Data curation, Formal analysis, Investigation, Methodology, Validation, Visualization, Writing – review and editing.

**Lydia J. Pung** Data curation, Formal analysis, Validation, Visualization, Writing – review and editing.

**Lina Gao** Data curation, Formal analysis, Investigation, Methodology, Validation, Writing – review and editing.

**Zachary J. Streblow** Data curation, Formal analysis, Investigation, Methodology, Validation, Visualization, Writing – review and editing.

**Brayden Graves** Data curation, Formal analysis, Writing – review and editing.

**Magdalene M. Streblow** Data curation, Formal analysis, Writing – review and editing.

**Aaron M. Barber-Axthelm** Data curation, Investigation, Methodology, Writing – review & editing.

**Gavin Zilverberg** Data curation, Investigation, Methodology, Writing – review & editing.

**Margaret Terry** Data curation, Investigation, Methodology, Writing – review & editing.

**Suzanne S. Fei** Data curation, Formal analysis, Investigation, Methodology, Validation, Visualization, Writing – review and editing.

**Glenn Hogan** Data curation, Formal analysis, Methodology, Validation, Visualization, Writing – review and editing.

**David C. Schultz** Data curation, Formal analysis, Methodology, Validation, Visualization, Writing – review and editing.

**Sara Cherry** Conceptualization, Data curation, Formal analysis, Investigation, Methodology, Validation, Visualization, Writing – review and editing.

**Michael K. Axthelm** Conceptualization, Data curation, Formal analysis, Investigation, Methodology, Validation, Visualization, Writing – review and editing.

**Caralyn S. Labriola** Conceptualization, Data curation, Formal analysis, Investigation, Methodology, Validation, Visualization, Writing – original draft.

**Mark Heise** Conceptualization, Data curation, Formal analysis, Investigation, Methodology, Validation, Visualization, Writing – review and editing.

**Daniel N. Streblow** Conceptualization, Data curation, Formal analysis, Investigation, Methodology, Validation, Visualization, Supervision, Writing – original draft.

## Institutional Review Board Statement

The study protocol was approved by the Institutional Animal Care and Use Committee of Oregon Health & Science University (protocol code: 0993; expiration date: 12 April 2026).

## Data Availability Statement

The original contributions presented in the study are included in the article/supplementary material; Raw data of transcriptome analysis will be submitted to the SRA (pending processing). Further inquiries can be directed to the corresponding author.

## Acknowledgments

The authors acknowledge the Integrated Pathology Core at the ONPRC for the preparation of histologic slides and the UPenn High-throughput Screening core, RRID: SCD_022379, for assistance with the Fc reporter assays. We would also like to thank the veterinarians and husbandry staff, and the Infectious Disease Resource at the ONPRC who provided excellent care for the animals used in this study.

